# Isolation and whole genome characterization of antagonistic *Paenibacillus polymyxa* 188 and its biocontrol potential against several fungi

**DOI:** 10.1101/2023.03.22.533881

**Authors:** Sra-Yh Shih, Ker-Rui Chou, Hung-Yi Wu, HsinYuan Tsai

**Affiliations:** Department of Marine Biotechnology and Resources, National Sun Yat-Sen University, Kaohsiung City, Taiwan

## Abstract

Discovery of new antifungal compounds and biocontrol agents is important due to the emergence of drug-resistant fungi and new fungal pathogens. The aim of this study was to isolate marine bacteria that are able to produce metabolites with antifungal properties. In this study, we isolated a *P. polymyxa,* named as 188, from a marine sediment sample and evaluated its antifungal ability. The results indicated that the bacterium showed excellent antifungal activity against many pathogenic fungi of plants and humans. The antifungal compounds produced by *P. polymyxa* 188 were extracted and analyzed using MALDI-TOF/MS. The complete genome sequence and biosynthetic gene clusters were characterized, and further to compare the genomes of *P. polymyxa* 188 with other strains. Various CAZymes were identified in *P. polymyxa* 188. Five antibiotic gene clusters including paenilan, paenibacillin, fusaricidin, polymyxin, and tridecaptin can be found in *P. polymyxa* strains, but gene clusters of paenilan, paenibacillin, and polymyxin were absent in *P. polymyxa* 188. Our findings provided detail genetic information about *P. polymyxa* 188 and suggested that *P. polymyxa* 188 is the potential agent of biocontrol and disease management.

## Introduction

Chemical fertilizers and pesticides have been widely utilized in plant agriculture to control pests and diseases and improve the quality and yield of commercial plants. However, overuse of these chemicals has emerged as a serious problem for environmental contamination and human health hazards (Sindhu et al., 2016). The long-term usage of chemicals leads to the development and spread of antimicrobial resistance (AMR) and reduces microbial diversity in soil (Chen et al., 2019; Miller et al., 2022). Therefore, using safe and eco-friendly biocontrol agents as an alternative to conventional farming strategies is urgently needed (Jiang et al., 2022). Biocontrol agents refer to living organisms; they fight against plant diseases and pests through direct antagonistic effects as well as the indirect stimulation of plant resistance. Biocontrol agent has been extensively studied in many microorganisms such as *Bacillus subtilis* (Kloepper et al., 2004), *Pseudomonas fluorescens* (Nagarajkumar et al., 2004), *Serratia plymuthica* (Kamensky et al., 2003), and *Paenibacillus polymyxa,* all have been demonstrated the antagonistic activity against plant pathogens.

*P. polymyxa*, a Gram-positive and plant growth-promoting rhizobacterium (PGPR), is found in diverse habitats including soil rhizospheres, plant roots, marine sediments, and even the gut of insects (Guemouri-Athmani et al., 2000; von derWeid et al., 2000; Lal and Tabacchioni, 2009; Soni et al., 2021). Many strains of *P. polymyxa* are known to have positive effects on plants through multiple mechanisms including biosynthesis of plant growth hormones, nitrogen fixation, inorganic phosphate solubilization, and improvement of abiotic and biotic stress tolerance (Bohlool et al., 1992; Weselowski et al., 2016a; Backer et al., 2018). They have been applied to various crops, such as strawberry, cucumber, tomato, and tobacco (Hao and Chen, 2017; Liu et al., 2020; Tsai et al., 2021; Du et al., 2022), and used for biological control of pathogenic fungi or bacteria (Dijksterhuis et al. 1999; Timmusk and Wagner 1999; Helbig 2001; Beatty and Jensen 2002; Yang et al. 2002; Ryu et al. 2006; Raza et al. 2008). *P. polymyxa* strains have been reported to show broad-spectrum antifungal activity against a variety of species, such as *Fusarium oxysporum, Leptosphaeria maculans, Coniella diplodiella, Aspergillus niger, Phytophora palmivora* and *Phytim aphanidermatum* (Beatty and Jensen 2002; Haggag 2007; Han et al. 2015; Raza et al. 2015; Timmusk et al. 2003), as well as against plant pathogenic bacteria such as *Pseudomonas syringae* (Weselowski et al., 2016b) and *Ralstonia solanacearum* (Yi et al., 2019).

Several types of antibiotics including fusaricidin, paenilan, polymyxin, tridecaptin, and paenibacillin can be produced by *P. polymyxa* (Luo et al., 2018; Jeong et al., 2019). Of these, fusaricidins are the common antifungal secondary metabolites (SMs) produced by *P. polymyxa,* which are composed of a ring peptide with six amino acids and a 15-guanidino-3-hydroxypentadecanoic acid (GHPD) as a side chain (Kajimura and Kaneda, 1996; Vater et al., 2015). Fusaricidins display high antimicrobial activity against fungi and Gram-positive bacteria, but shows limited efficacy against Gram-negative bacteria (Cochrane and Vederas, 2016). The antibiotic can inhibit the germination of *Fusarium* spores and interfere with the integrity of hyphal membranes (Li and Chen, 2019). At least 40 kinds of fusaricidins and its analogs have been reported from different *P. polymyxa* strains such as fusaricidin A, fusaricidin A1, fusaricidin B, fusaricidin B1, fusaricidin C, fusaricidin D, fusaricidin E, fusaricidin F and LI-F03 to LI-F08 (Choi et al., 2007; Vater et al., 2015; Qiu et al., 2019). The fusaricidin biosynthetic gene cluster (BGC) contains eight genes including *fusG, fusF, fusE, fusD, fusC, fusB, fusA,* and *fusTE.* These genes except for *fusTE* are associated with fusaricidins production (Li and Chen, 2019), and previous studies showed that fusaricidin production is controlled and regulated by the KinB-Spo0A-AbrB pathway in *P. polymyxa* WLY78 (Li et al., 2021).

The genomic comparison and transcriptome analysis are beneficial for understanding the genetic biocontrol mechanisms in *P. polymyxa* strains (Xie et al., 2016; Liu et al., 2021; Soni et al., 2021; Du et al., 2022). To date, ninety-five *P. polymyxa* genome assemblies have been deposited at NCBI database (https://www.ncbi.nlm.nih.gov/, date accessed 28/Dec/2022). In this study, a novel *P. polymyxa* 188 was isolated from marine sediment showing significant antifungal activity. However, its whole genome was uncharacterized and the biocontrol mechanism remains unclear. The aims of this study were to (1) identify the antifungal metabolites; (2) evaluate the antagonistic activity of *P. polymyxa* 188 against pathogenic fungi; (3) characterize the *P. polymyxa* 188 genome architecture and perform a comparative genomic analysis across different *P. polymyxa*strains; (4) mine the potential SMs in *P*. *polymyxa* strains.

## Methods and Materials

### Bacterial isolation

*P. polymyxa* 188 was isolated from a marine sediment sample in western Taiwan. The sediment sample was collected in a 50-mL sterile centrifuge tube and delivered to the laboratory within 24 hours. The sample was diluted with 2.5% NaCl solution and plated on mitis-salivarius agar at 30°C for 48 hours. Restreaking was performed to obtain a pure culture. The pure bacterial isolate was identified by 16S rRNA gene sequencing and preserved in 20% glycerol at −80°C. The strain was routinely cultured in 523 broth (1% sucrose, 0.8% casein hydrolysate, 0.4% yeast extract, 0.2% K2HPO4, 0.03% MgSO4, pH 6.9)(Kado and Heskett, 1970) at 30°C at 200 rpm.

### Antagonism assay

The antagonistic ability of the isolate was evaluated by a dual-culture assay against tested fungi including *Aspergillus terreus, Cochliobolus* sp., *Fusarium oxysporum f*. sp. *cubense, Fusarium tricinctum, Fusarium oxysporum, Microsphaeropsis arundinis, Pestalotiopsis clavispora, Penicillium oxalicum* and *Talaromyces pinophilus.* Dual-culture assays were performed in 6 cm diameter Petri plates containing potato dextrose agar (PDA) medium. The fungal mycelium was placed at 2.5 cm from the edge of the plate and a 5 cm long band of *P. polymyxa* 188 was streaked on PDA at a distance of 0.5 cm from the fungal mycelium. The PDA plate inoculated with fungal mycelium only was used as a control. After 72 hours of incubation, the results of the antagonist assay were observed and photographed. The percentage of inhibition of fungal growth was measured by the following formula described by Yassin et al. (2021).

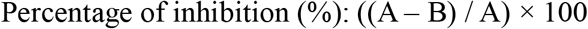

where A is diameter of tested fungus on control plate and B is diameter of tested fungus on treated plate.

### Extraction and identification of antifungal activity fractions

To extract the antifungal compounds, the pellet of *P. polymyxa* 188 was extracted by methanol (added 3 mL of methanol per 100 μL pellet) for 16 hours. After centrifugation, the crude extract was air-dried and then dissolved in 1 mL of methanol. The crude extract was further fractionated by Normal phase chromatography (NP). One mL crude extract was loaded onto the column and eluted with ethyl acetate:methanol (40:60, v:v) in 0.1% trifluoroacetic acid solution at a flow rate of 1 mL/min. The fractions were air-dried and dissolved in 500 μL methanol. The active fraction was detected by using matrix-assisted laser desorption ionization-time of flight mass spectrometry (MALDI-TOF MS). Mass spectra were recorded on a MALDI-TOF MS (AutoFlex III, Bruker Daltonics, Germany) equipped with Nd-YAG laser (355nm) for desorption and ionization. The sample mixture was mixed with an equal volume of matrix solution (α-Cyano-4-hydroxycinnamic acid), spotted on the target, air-dried and measured. Spectra were obtained by positive ion detection and reflector mode. Monoisotopic masses were recorded.

### Genome sequencing, assembly, and annotation

Genomic DNA of *P. polymyxa* 188 was extracted using Phenol Chloroform Isoamyl alcohol method (Aamir, 2015). The quality of isolated DNA was analyzed by 1% agarose gel electrophoresis. The whole genome sequence of *P*. *polymyxa* 188 was performed using PacBio RS II platform (Pacific Biosciences, USA). The sequence data were assembled and polished using hifiasm v0.15.3 and GCpp v2.0.2, respectively. The assembled genome quality was evaluated using QUAST v5.0.2 (Gurevich et al., 2013). The assembled genome was annotated using Prokka v1.14.6 (Seemann, 2014). The graphical circular map was generated using CGview comparison tool. Tandem repeats and insertion sequences were identified using tandem repeats finder (Benson, 1999) and IS Finder (Siguier et al., 2006), respectively. Genomic islands (GIs) and the prophages were predicted with IslandViewer 4 (Zhang et al., 2000) and PHASTER (https://phaster.ca/), respectively. The predicted proteins were assigned by the Cluster of orthologous groups (COG) using eggNOG-mapper (Cantalapiedra et al., 2021).

### Phylogenetic analysis and genome comparison

Genome assemblies of 90 *P*. *polymyxa* strains were retrieved from the NCBI (accessed on June 2022). The phylogenetic tree of *P*. *polymyxa* strains was constructed using PYANI v.0.2.11 based on BLAST+ algorithm (ANIb) calculation (Pritchard et al., 2016) and iTOL web server (Letunic and Bork, 2021), and *P*. *polymyxa,* which have contig < 3 and average nucleotide identity (ANI) > 92% verse *P*. *polymyxa* 188, were annotated using the Prokaryotic Genome Annotation System (Prokka) v1.12 (Seemann, 2014). *In silico* DNA-DNA Hybridization (DDH) across *P*. *polymyxa* was calculated using Genome to Genome Distance Calculator 3.0 (http://ggdc.dsmz.de/home.php). Carbohydrate-active enzymes (CAZymes) including glycoside hydrolases (GHs), glycosyltransferases (GTs), polysaccharide lyases (PLs), carbohydrate esterases (CEs), auxiliary activities (AAs), and carbohydrate-binding modules (CBMs), were identified using dbCAN2 meta server (Zhang et al., 2018) (accessed on November 2021) with HAMMER, DIAMOND, and dbCAN_sub. Secondary metabolite gene clusters were predicted using the antiSMASH v. 6.0.1 (bacterial version) (Blin et al., 2021) with the default mode.

## Results

### Antagonist assay of *P. polymyxa* 188 against nine fungi

To evaluate the antagonistic activity of *P. polymyxa* 188 against nine fungi, a dual culture assay was conducted. The results showed that *P. polymyxa* 188 exhibited antifungal activities against all of the tested fungi (Figure 1). Specifically, *P. polymyxa* 188 almost completely inhibited the growth of *M. arundinis* (98%) and with inhibition of *F. tricinctum* (96%), *P. clavispora* (92%), *Fusarium oxysporum* (92%), *F. oxysporum f.* sp. *Cubense* (90%), and *P. oxalicum* (85%). Additionally, *P. polymyxa* 188 inhibited the growth of *A. terreus, Cochliobolus* sp., and *T. pinophilus* by approximately 80%, 68%, and 63%, respectively.

**Figure 1.**
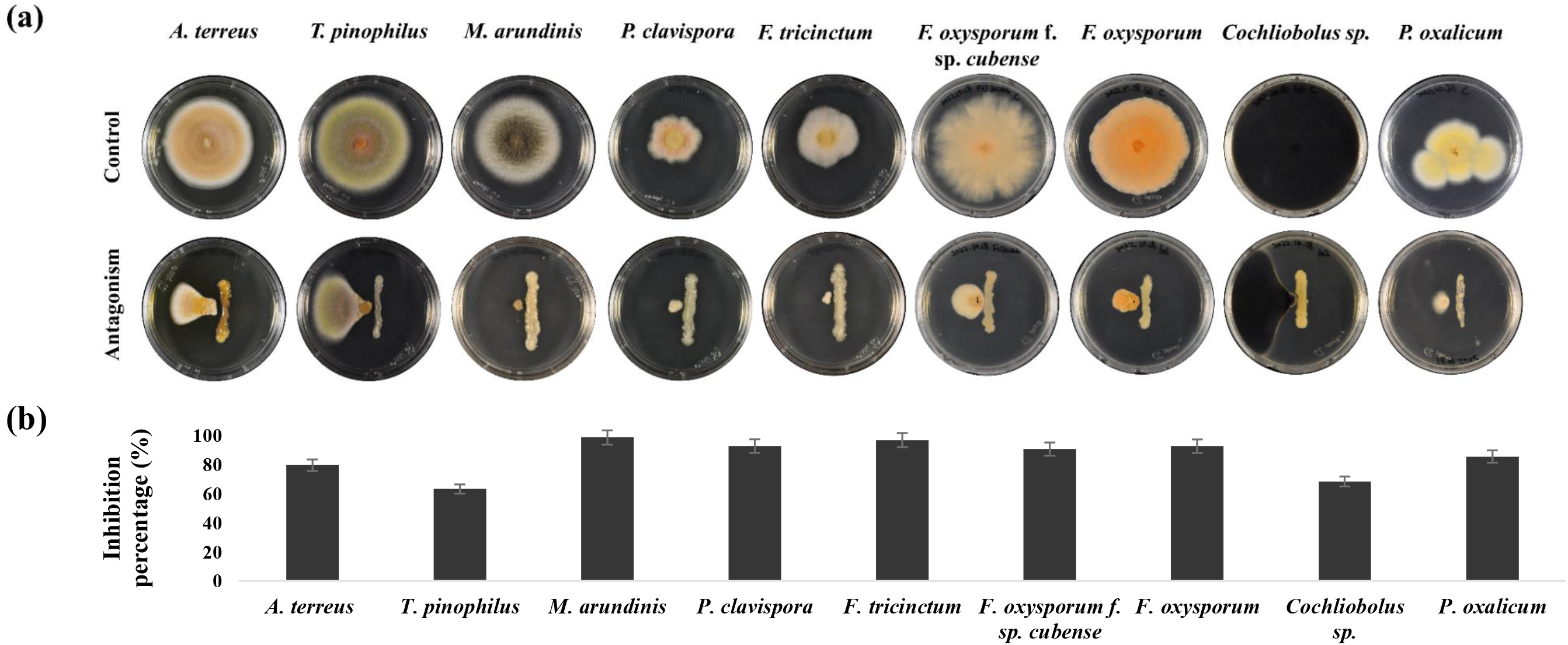
In vitro antagonistic activity of *P. polymyxa* 188 against different fungal pathogens. (a) For each assay, a single culture of the tested fungus was used as the control. In dual culture assays, the tested fungus was plated on the left, and the *P. polymyxa* 188 was plated on the right. The tested fungi were *Aspergillus terreus, Talaromyces pinophilus, Microsphaeropsis arundinis, Pestalotiopsis clavispora, Fusarium tricinctum, Fusarium oxysporum f*. sp. *cubense, Fusarium oxysporum, Cochliobolus* sp., and *Penicillium oxalicum.* (b) The percentages of inhibition of fungal growth were measured in triplicate. Error bars represented standard error of the means.

### Extraction and identification of antifungal compound produced by *P. polymyxa* 188

The extraction of *P. polymyxa* 188 was made using methanol. The fraction mixture analyzed with MALDI-TOF showed multiple peaks at m/z 860-1040. The mass spectrum was depicted in Figure 2 and the mass data was summarized in Table 1. The spectrum results revealed that the mass peaks were attributed to fusaricidin, including fusaricidin A (m/z 883), fusaricidin B (m/z 897), fusaricidin C (m/z 947), and fusaricidin D (m/z 961). The minor mass peaks were related to LI-F type compounds, such as LI-F04b, LI-F05a, LI-F06a, LI-F05b, LI-F06b, LI-F03a, LI-F03b, LI-F07b, and LI-F08b. The mass peaks, including m/z 905, 919, 921, and 935 were reported by Vater et al. (2015), and m/z 963, 967, 969, 983, 985, and 999 were reported by Vater et al. (2016) as fusaricidin-type compounds.

**Figure 2.**
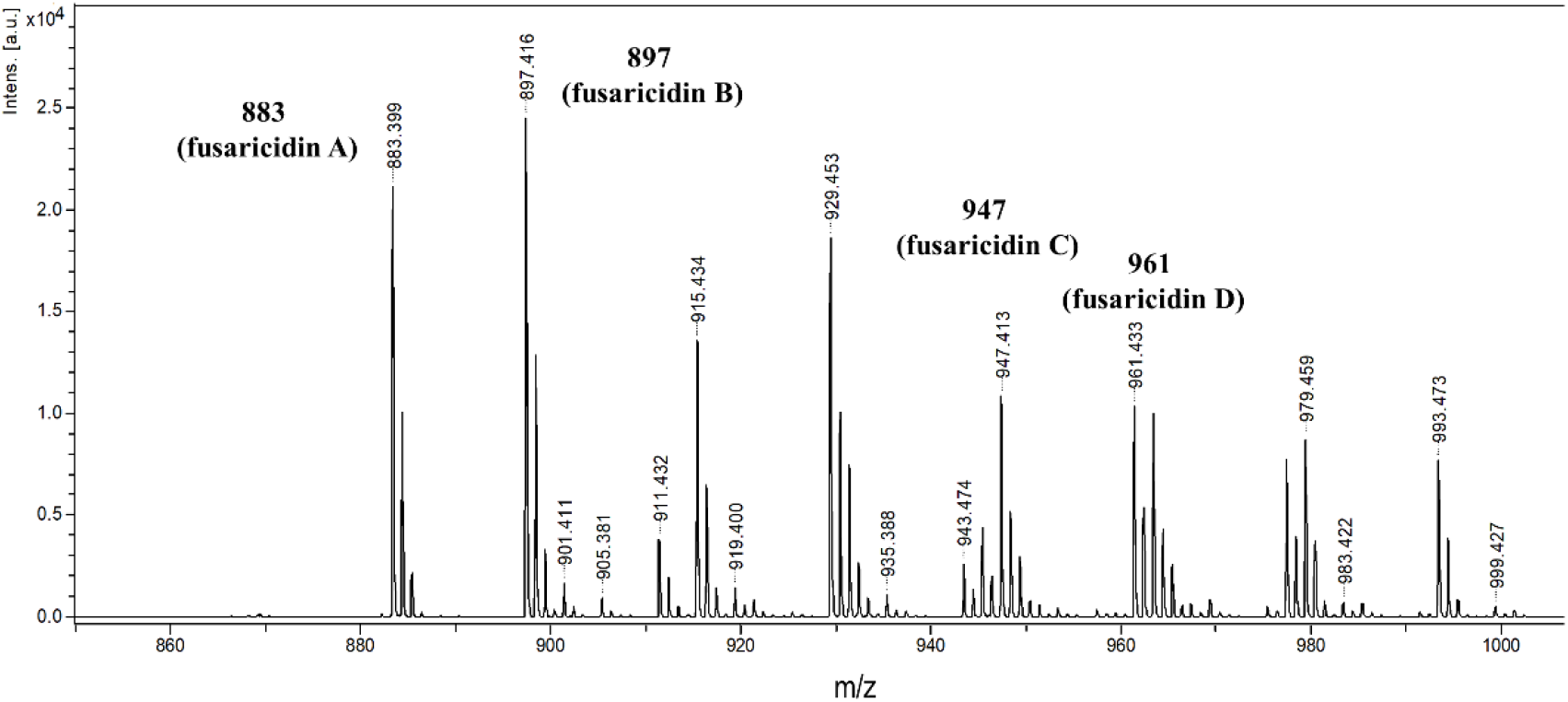
MALDI-TOF analysis of antifungal substances obtained from *P. polymyxa* 188. The mass spectrum of fusaricidin-type compounds were identified (m/z: 883, 897, 905, 911, 919, 935, 947, 961, 983, and 999).

**Table 1.** Fusaricidin-type compounds were detected in active antifungal fraction from *P. polymyxa* 188 by MALDI-TOF Mass spectrometry.

| **[M+H]<sup>+</sup>** | **Fusaricidin** | **References** |
| --- | --- | --- |
| m/z |  |  |
| 883 / 905 / 921 | Fusaricidin A | Kajimura and Kaneda 1996 |
| 897 / 919 / 935 | Fusaricidin B, LI-F04b<br>LI-F05a, LI-F06a | Kajimura and Kaneda 1997; Kurusu et al. 1987; Nomura et al. 2000 |
| 911 | LI-F05b, LI-F06b | Kurusu et al. 1987; Nomura et al. 2000 |
| 947 / 969 / 985 | Fusaricidin C, LI-F03a | Kajimura and Kaneda 1997 |
| 961 / 983 / 999 | Fusaricidin D, LI-F03b | Kajimura and Kaneda 1997 |
| 963 | LI-F08b | Kurusu et al. 1987; Kuroda et al. 2000 |
| 967 | LI-F07b | Kurusu et al. 1987; Kuroda et al. 2000 |

### Genomic features of *P*. *polymyxa* 188

The complete genome of *P*. *polymyxa* 188 generated 5,542,145 bp with one contig (~296× coverage). The GC content was 45.5%. There were 5,119 genes identified including 4,912 protein-coding genes and 207 RNA genes (110 tRNA, 40 rRNA, and 56 misc RNA) (Table 2). The graphical circular map was shown in Figure 3. Cluster of orthologous groups of proteins (COG) analysis categorized 3,774 genes, and the most frequent category (44.3%) was “metabolism” (Table 2; Figure 4). The full tandem repeats, genomic islands (GIs), Phaster, and insertion sequences predicted based on *P*. *polymyxa* 188 were listed in Table S1-S4.

**Figure 3.**
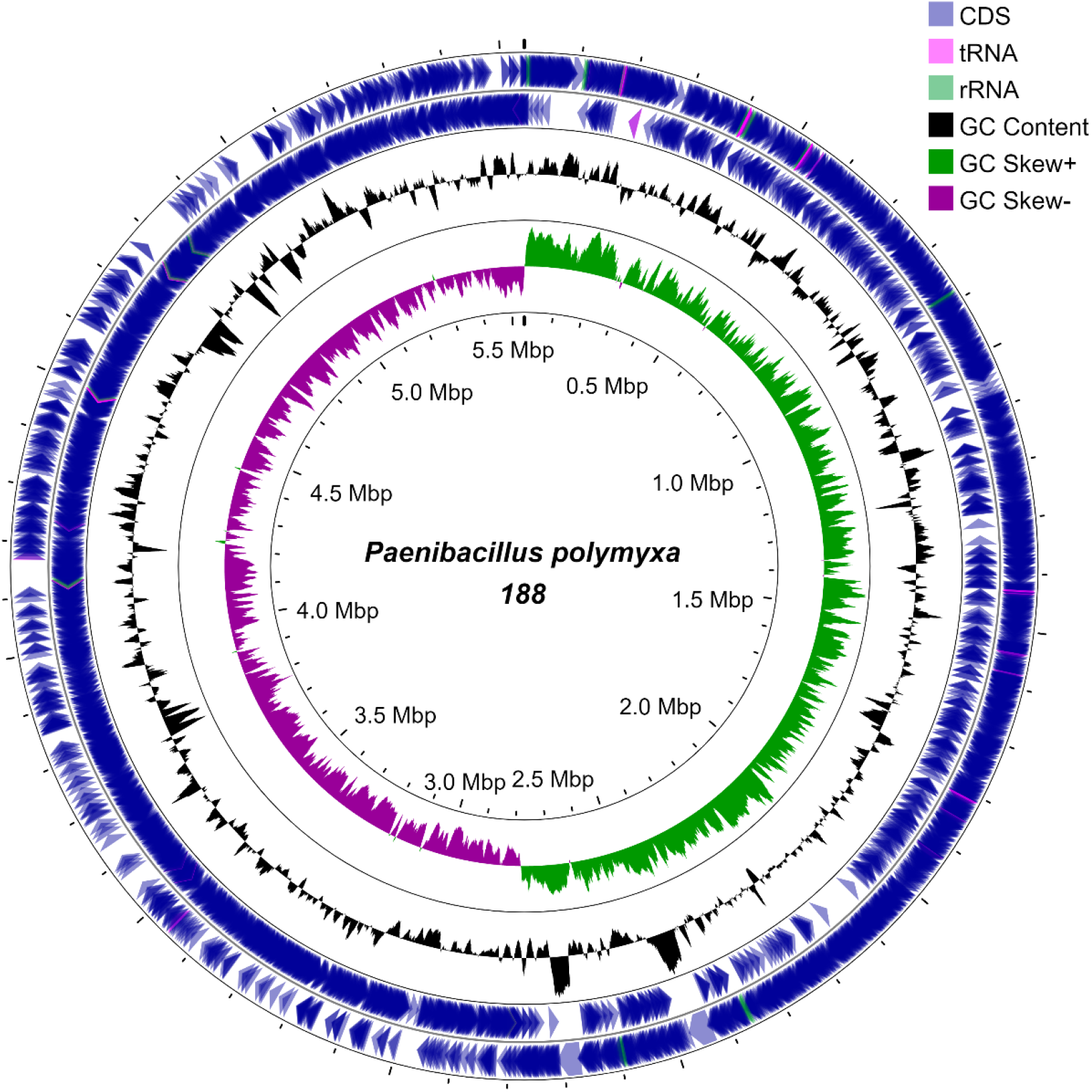
Graphical circular map of *P*. *polymyxa* 188 chromosome. From outside to inside rings: Ring 1 and Ring 2 represented forward and reverse CDS with tRNA and rRNA. Ring 3 showed the GC content, and Ring 4 showed the GC skew (green and purple).

**Figure 4.**
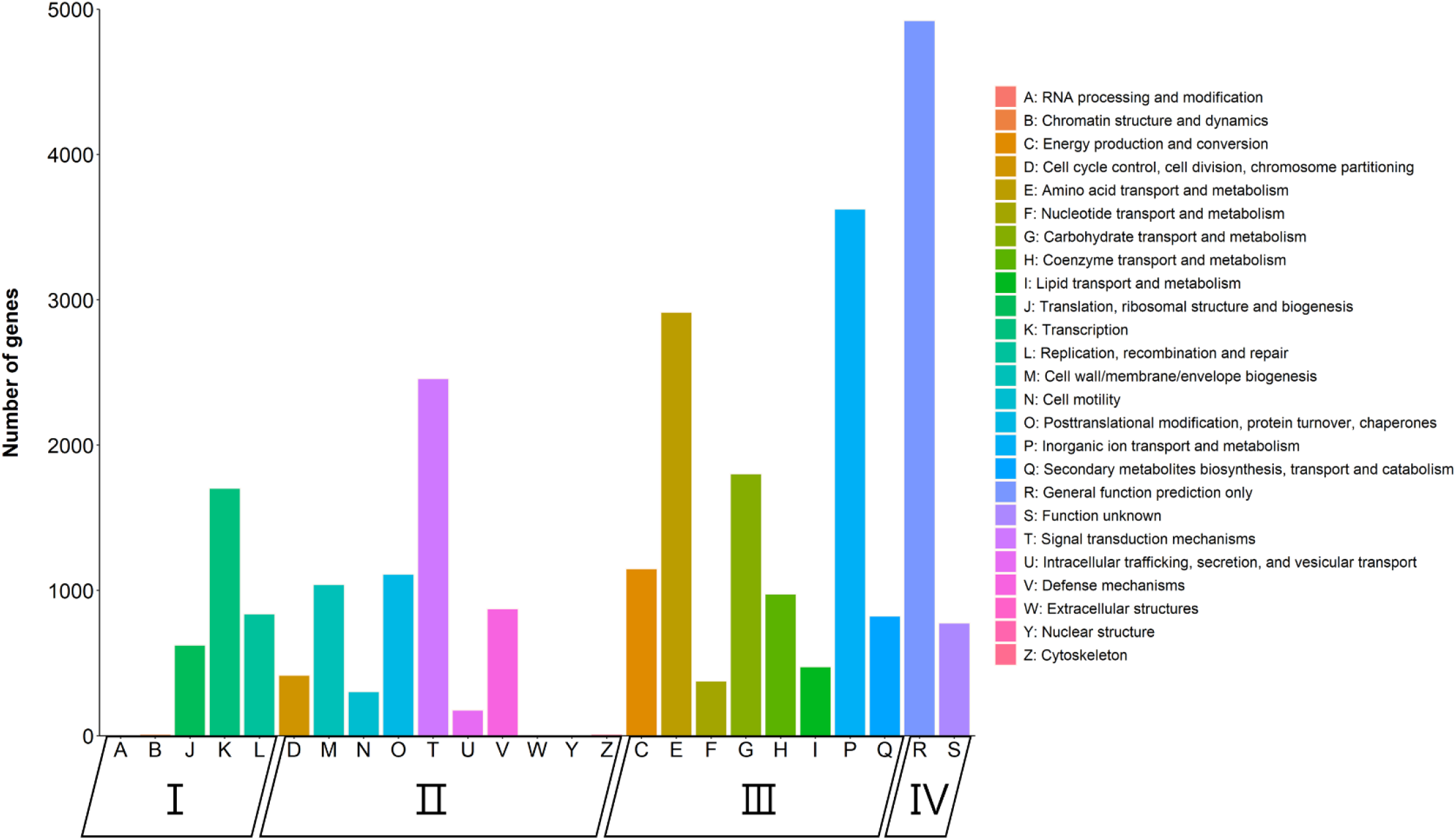
COG functional categories in *P. polymyxa* 188. COG function classification were assigned to four major groups including classification information storage and processing (I), cellular processes and signaling (II), metabolism (III), and function poorly characterized proteins (IV). The X-axis represents gene functional category and the Y-axis represents the number of gene.

**Table 2.** General features of the genome of the *P. polymyxa* strain 188.

| Features | Value |
| --- | --- |
| Genome size (bp) | 5,542,145 |
| Genome coverage | 296× |
| Contig numbers | 1 |
| GC content (%) | 45.5 |
| Total number of genes | 5,119 |
| Protein-coding genes (CDSs) | 4,912 |
| tRNA genes | 110 |
| misc RNA | 56 |
| rRNA genes | 40 |
| tmRNA genes | 1 |
| COG annotated genes | 3,774 |
| Predicted CAZymes proteins | 238 |
| Secondary metabolite BGCs | 15 |
| Tandem repeats | 115 |
| Genomic islands | 22 |
| CRISPR | 1 |

### Phylogenetic relationship and comparative genomics of *P*. *polymyxa*

Twenty-two *P*. *polymyxa* strains were compared with *P*. *polymyxa* 188, showing the average genome size of the *P*.*polymyxa* strains was about 5.9 Mb and the average GC content was 45.4% (Table 3). The phylogenetic tree was divided into three major clades and *P*. *polymyxa* 188 was closely related to *P*. *polymyxa* ZF197 with highly ANI value (97.3 %) and DDH value (87.2 %) (Figure 5).

**Figure 5.**
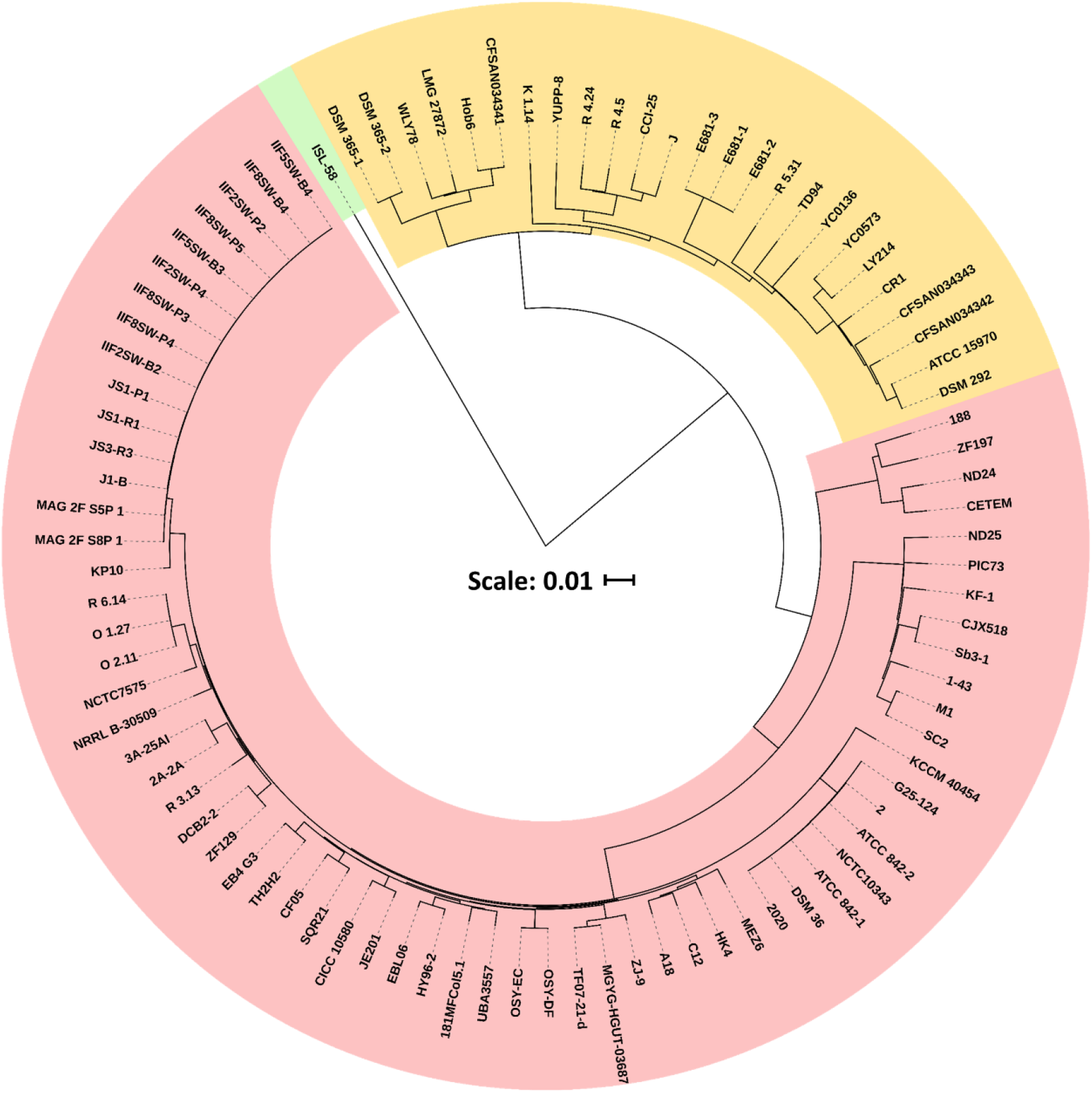
Phylogenetic tree of *P. polymyxa* stains. The phylogenetic tree based on ANI values was constructed using PYANI v.0.2.11 based on BLAST+ program and iTOL web server. The scale bar represented the sequence divergence.

**Table 3.** Comparisons of genome characteristic features of *P. polymyxa* stains.

| Strains | GenBank<br>assembly accession | Contig N50 | Chromosome<br>Size (bp) | CDS (n) | GC% | Scaffolds | ANI similarity versus<br>*P. polymyxa* 188 (%) | DDH versus<br>*P. polymyxa* 188 (%) |
| --- | --- | --- | --- | --- | --- | --- | --- | --- |
| 188 |  | 5,542,145 | 5,542,145 | 4,912 | 45.5 | 1 | - | - |
| ZF197 | GCA_007858415.1 | 5,507,169 | 5,539,234 | 4,986 | 45.6 | 2 | 97.3 | 87.2 |
| CJX518 | GCA_014854715.1 | 5,690,411 | 5,690,411 | 4,959 | 45.4 | 1 | 93.0 | 80.4 |
| Sb3-1 | GCA_000819665.1 | 5,597,590 | 5,829,236 | 5,018 | 45.4 | 3 | 93.0 | 78.4 |
| SC2 | GCA_000164985.2 | 5,728,392 | 6,238,510 | 5,707 | 44.6 | 2 | 92.9 | 71.6 |
| M1 | GCA_000237325.1 | 5,864,546 | 6,231,122 | 5,591 | 44.8 | 2 | 92.9 | 72.1 |
| ATCC_842-1 | GCA_022811565.1 | 5,921,998 | 5,967,522 | 5,390 | 45.1 | 2 | 92.5 | 74.5 |
| O_2.11 | GCA_023586805.1 | 5,495,678 | 5,744,058 | 4,933 | 45.5 | 2 | 92.5 | 77.8 |
| C12 | GCA_022649565.1 | 5,793,823 | 5,793,823 | 5,041 | 45.6 | 1 | 92.5 | 77.1 |
| NCTC10343 | GCA_900454525.1 | 5,933,430 | 5,984,949 | 5,405 | 45.1 | 2 | 92.5 | 74.5 |
| ZJ-9 | GCA_022492955.1 | 5,755,887 | 5,755,887 | 5,006 | 45.7 | 1 | 92.5 | 77.2 |
| ZF129 | GCA_006274405.1 | 5,703,931 | 5,820,553 | 5,072 | 45.4 | 3 | 92.5 | 75.9 |
| R_3.13 | GCA_023586645.1 | 5,730,787 | 5,743,686 | 4,996 | 45.6 | 2 | 92.5 | 77.6 |
| HY96-2 | GCA_002893885.1 | 5,745,779 | 5,745,779 | 4,936 | 45.6 | 1 | 92.5 | 78.3 |
| O_1.27 | GCA_023586745.1 | 5,493,315 | 5,797,190 | 4,935 | 45.6 | 3 | 92.5 | 77.9 |

Table 3. (continued) Comparisons of genome characteristic features of *P. polymyxa* stains.
| Strains | GenBank<br>assembly accession | Contig N50 | Chromosome<br>Size (bp) | CDS (n) | GC% | Scaffolds | ANI similarity versus<br>*P. polymyxa* 188 (%) | DDH versus<br>*P. polymyxa* 188 (%) |
| --- | --- | --- | --- | --- | --- | --- | --- | --- |
| 2020 | GCA_015710815.1 | 5,916,219 | 5,961,740 | 5,497 | 45.1 | 2 | 92.4 | 74.4 |
| DSM_36 | GCA_015710975.1 | 5,916,392 | 5,961,915 | 5,447 | 45.1 | 2 | 92.4 | 74.4 |
| MEZ6 | GCA_023374295.1 | 5,769,542 | 5,769,542 | 4,910 | 45.6 | 1 | 92.4 | 77.3 |
| R_6.14 | GCA_023586785.1 | 5,745,819 | 5,745,819 | 4,915 | 45.5 | 1 | 92.4 | 77.8 |
| SQR21 | GCA_000597985.1 | 5,828,436 | 5,828,436 | 5,024 | 45.7 | 1 | 92.4 | 77.4 |
| CF05 | GCA_000785455.1 | 3,188,772 | 5,762,608 | 4,919 | 45.5 | 1 | 92.4 | 77.6 |
| JE201 | GCA_019852195.1 | 5,610,112 | 6,167,064 | 5,358 | 45.3 | 3 | 92.4 | 73.6 |
| EB4_G3 | GCA_020181695.1 | 5,739,603 | 5,739,603 | 4,944 | 45.8 | 1 | 92.4 | 77.5 |

### CAZyme and antibiotic gene clusters

238 proteins were assigned to CAZyme families including 115 glycosyl hydrolases (GHs), 73 glycosyltransferases (GTs), 28 carbohydrate esterases (CEs), 4 auxiliary activities (AAs), and 10 polysaccharide lyase (PLs), 31 carbon-binding domain (CBMs). *P*. *polymyxa* 188 had the highest number of GTs across different *P*. *polymyxa* strains (Table S5; Figure 6). The comparison of five antibiotic gene clusters (paenilan, paenibacillin, fusaricidin, polymyxin, and tridecaptin) showed that *P*. *polymyxa* 188 and ZF197 were lack of paenilan, paenibacillin, and polymyxin. In addition, Fusaricidin biosynthetic gene cluster (BGC) was found in all *P*. *polymyxa* strains, but paenibacillin BGC only existed in some strains of *P. polymyxa* including ATCC 842-1, C12, NCTC10343, ZF129, R3.13, EB4 G3, 2020, and DSM 36 (Figure 7).

**Figure 6.**
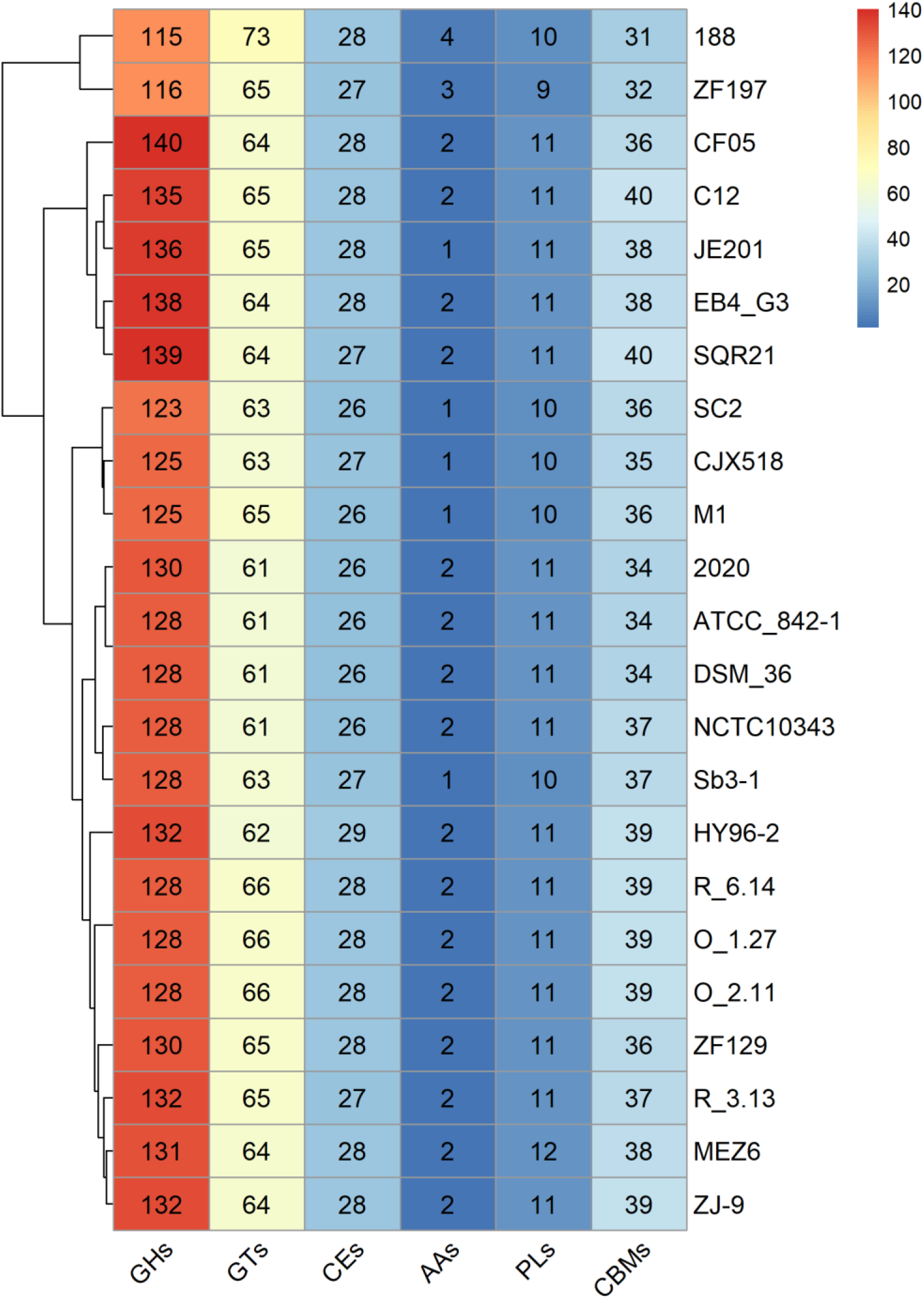
Predicted numbers of CAZyme in *P. polymyxa* stains. X-axis: CAZyme. Y-axis: Strains.

**Figure 7.**
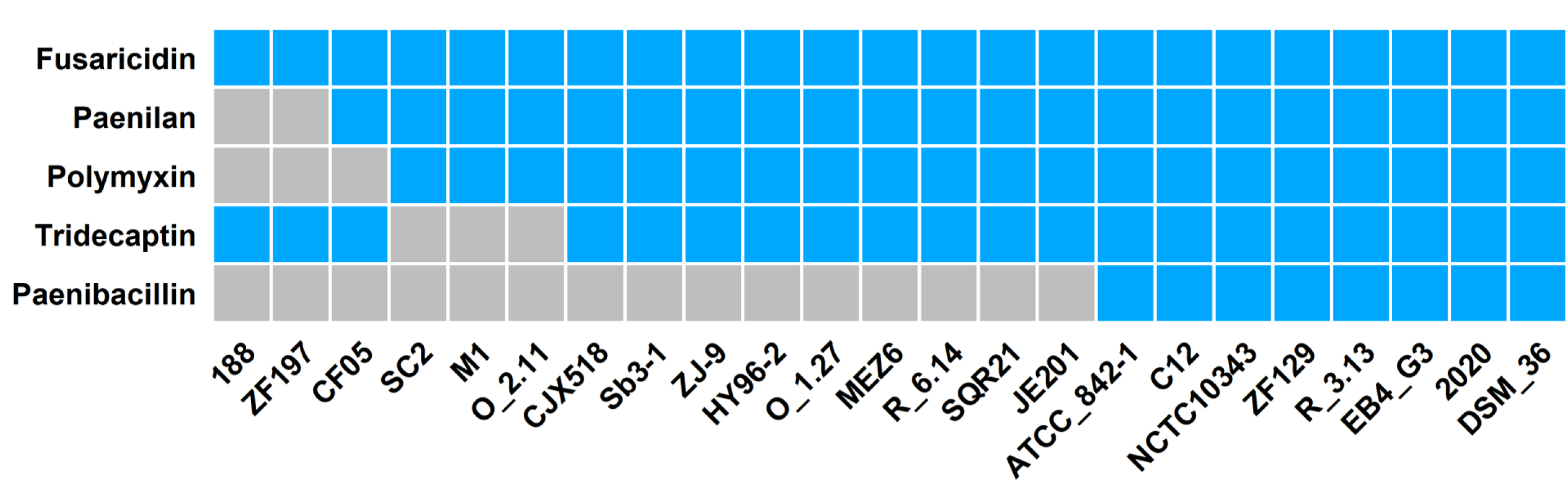
Secondary metabolites (SMs) of *P. polymyxa* strains. X-axis: Strains. Y-axis: SMs.

## Discussion

In this study, we isolated a bacterium from marine sediment exhibiting high antifungal activity. The 16S rRNA sequence analysis revealed that the bacterial isolate named *P. polymyxa* 188 belonged to the genus Paenibacillus, showing the highest sequence similarity to *P. polymyxa* ZF197 (97.31%). Unlike *P. polymyxa* ZF197, which was obtained from potato rhizospheric soil, *P. polymyxa* 188 was isolated from marine sediment. Studies reported that *P. polymyxa* mostly found in soil and plant rhizosphere (Lal and Tabacchioni 2009; Padda et al. 2017). One common feature of *P. polymyxa* is its ability to inhibit a variety of fungi. Therefore, we conducted antagonism assays on *P. polymyxa* 188 against nine fungi to investigate its antifungal ability. *P. polymyxa* 188 was found to significantly inhibit the mycelial growth of all tested fungi, including six crop pathogenic fungi: *F. tricinctum* (Wang et al., 2022), *P. clavispora* (Valencia et al., 2011), *Fusarium oxysporum* (Gordon, 2017), *F. oxysporum f.* sp. *Cubense* (Magdama et al., 2020), and three human pathogenic fungi: *A. terreus* (Iwen et al., 1998), *P oxalicum* (Chowdhary et al., 2014) and *M. arundinis* (Reppas et al., 2015).

The extraction techniques were applied to determine bioactive compounds with antifungal properties in *P. polymyxa* 188. Beatty and Jensen (2002) described that antifungal compounds could not be extracted from the supernatant fraction, but from the harvested pellet using methanol as the solvent. Accordingly, we extracted antifungal compounds from the cell pellet using methanol. The bioactive fraction of *P. polymyxa* 188 was further identified using MALDI-TOF analysis and confirmed the presence of fusaricidins and LI-F compounds. Fusaricidins are the main antifungal compounds produced by *P. polymyxa*. Four types of fusaricidins (fusaricidin A-D) and nine kinds of LI-F compounds, including LI-F04b, LI-F05a, LI-F06a, LI-F05b, LI-F06b, LI-F03a, LI-F03b, LI-F07b, and LI-F08b were observed in *P. polymyxa* 188 extracts. Fusaricidins A, B, C, and D can effectively inhibit the growth of plant pathogenic fungi such as *Fusarium oxysporum, Aspergillus niger, Aspergillus oryzae,* and *Penicillum thomii*, and among these fusaricidin antibiotics, fusaricidin A showed outstanding antimicrobial activity against a range of clinical fungi and Gram-positive bacteria (Siepe et al., 2020). A study (Raza et al., 2009) demonstrated that fusaricidin A to D produced by *P. polymyxa* SQR-21 displayed antifungal activity against *Fusarium oxysporum* f. sp. *Nevium* (Raza et al., 2009). LI-F antibiotics produced by *P. polymyxa* L-1129 exhibited antifungal activity as well as activity against *Staphylococcus aureus* (Kuroda et al., 2000).

For comparative genomics studies, we compared the genomic sequences of *polymyxa*

188 with ninety *P*. *polymyxa* strains, and found twenty-two *P*. *polymyxa* strains had contig < 3 and high ANI similarity (> 92%) with the *P*. *polymyxa* 188. Based on ANI, *in silico* DDH, the results indicated that *P*. *polymyxa* 188 is highly similar to *P*. *polymyxa* ZF197 (97.3% ANI and 87.2%*in silico* DDH) (Table 3). *P*. *polymyxa* ZF197 is highly active against some plant-pathogenic fungi and bacteria, including *Verticillium dahlia, Fusarium oxysporum, Ralstonia solanacearum,* and *Pseudomonas syringae pv. Tomato* (Li et al., 2020). *P*. *polymyxa* strains are well-known for plant growth-promoting traits and produce several antimicrobial SMs, including paenilan, paenibacillin, fusaricidin, polymyxin, and tridecaptin, to control plant pathogens (Jeong et al., 2019; Soni et al., 2021). However, we found only two antimicrobial SMs, including fusaricidin and tridecaptin, in *P*. *polymyxa* 188 and ZF197. Fusaricidin SM was highly conserved across these strains, but paenibacillin SM only existed in *P*. *polymyxa* ATCC_842-1, C12, NCTC10343, ZF 129, R_3.13, EB4_G3, 2020, and DSM_36 (Figure 7). The gene clusters of fusaricidin and tridecaptin were first identified in *P*. *polymyxa* E681 (Jeong et al., 2019). Fusaricidin has activity against Gram-positive bacteria, plant pathogenic fungi, and oomycetes, whereas tridecaptin has activity against Gram-negative bacteria (Grady et al., 2016; Olishevska et al., 2019).

Additionally, *P*. *polymyxa* strains can produce various families of CAZymes, including GHs, GTs, CEs, AAs, PLs, and CBMs. These CAZymes give *P*. *polymyxa* good adaptability to different environments. *P*. *polymyxa* CF05, C12, JE201, EB4_G3, and SQR21 were comparatively high enrichment of GHs (135 to 140 genes), and *P*. *polymyxa* 188 had a higher number of GTs (73 genes) across *P*. *polymyxa* strains (Figure 6). According to Bohra et al. (2018), hemicellulolytic enzymes (GH2, GH10, GH11, GH16, GH26, GH30, GH31, GH36, GH43, GH51, GH74, and GH95 families) and cellulolytic enzymes (GH1, GH3, GH5, GH6, GH7, GH8 GH9, GH12, GH45, and GH48 families) were found in *Paenibacillus* spp. (López-Mondéjar et al., 2016; Bohra et al., 2018). *P*.*polymyxa* SQR21 had the highest number of GHs (38) associated with hemicellulose degradation, and *P*. *polymyxa* EB4 G3 and CF05 had the highest number of GHs (25) associated with cellulose degradation (Figure S1; Table S6). *P*. *polymyxa* strains, with the highest number of GHs involved in cellulolytic and hemicellulolytic deconstruction, were isolated from a fruit rhizosphere, a plant, and a tree (Shuqing et al., 2014; Mengying et al., 2015). These CAZymes can be utilized to produce biofuel, chemicals, food, and materials from plant biomass, and can replace chemical hydrolysis to degrade plant biomass in an environmentally friendly way.

## Conclusions

In the present study, we have shown that *P. polymyxa* 188 can inhibit various pathogenic fungi, and demonstrated the antifungal compounds are fusaricidin-type compounds. We characterized the genome architecture of *P. polymyxa* 188 and the genome analysis revealed that it is capable of producing various families of CAZymes, and two antibiotics, including fusaricidin and tridecaptin. Overall, our findings suggested that *P. polymyxa* 188 is a potential candidate for industrial use in CAZymes production as well as for biocontrol and disease management, due to its broad-spectrum antifungal activity. Furthermore, computational predictions based on whole genome sequence data provided further insights into the genetic characterization of *P. polymyxa* 188.

## Author contributions

SYS, KRC, HYW, and HYT conducted the experiments and prepared the manuscript. SYS and HYT conceived the investigation. All authors have read and agreed to the manuscript.

## Ethical statement

The experiments conducted on this study does not contain any studies with human participants or animals performed by any of the authors.

